# *Lactobacillus crispatus* S-layer proteins modulate innate immune response and inflammation in the lower female reproductive tract

**DOI:** 10.1101/2024.09.13.612838

**Authors:** Alexiane Decout, Ioannis Krasias, Lauren Roberts, Belen Gimeno Molina, Chloé Charenton, Daniel Brown Romero, Qiong Y. Tee, Julian R. Marchesi, Sherrianne Ng, Lynne Sykes, Phillip R Bennett, David A. MacIntyre

## Abstract

*Lactobacillus* species dominance of the vaginal microbiome is a hallmark of vaginal health. Pathogen displacement of vaginal lactobacilli drives innate immune activation and mucosal barrier disruption which increases the risks of STI acquisition and, in pregnancy, of preterm birth. Using cell reporter systems, we describe differential TLR mediated activation of the proinflammatory transcription factor NF-κB by vaginal pathogens and commensals. Vaginal *Lactobacillus* strains associated with optimal health were shown to selectively interact with anti-inflammatory innate immune receptors whereas species associated with suboptimal health including *L. iners* and *Gardnerella vaginalis* interacted with both pro- and anti-inflammatory receptors. Anti-inflammatory action of *L. crispatus* was regulated by surface layer protein (SLPs)-mediated shielding of TLR ligands and selective interaction with anti-inflammatory receptor, DC-SIGN. In pregnant women, cervicovaginal SLPs were predominately associated with *Lactobacillus-*enriched microbiota. These data offer new mechanistic insights into how vaginal microbiota modulate host immune response and influences risk of preterm birth.

## Introduction

Dominance of the vaginal microbiome by commensal *Lactobacillus* species, particularly *L. crispatus*, is widely associated with vaginal health. These species produce immunomodulatory and antimicrobial compounds that dampen inflammation and help to maintain the mucosal barrier thereby protecting against pathogen colonisation^1–5^. Conversely, *Lactobacillus-* depleted, high-diversity vaginal microbiota have been linked to mucosal inflammation^6,7^ and adverse health outcomes such as bacterial vaginosis, Sexually Transmitted Infections (STI) acquisition, miscarriage, and preterm birth^8–11^. Such non-optimal community compositions are often enriched with potentially pathogenic and BV-associated bacteria such as *Gardnerella vaginalis*, *Fannyhessea vaginae* and *Prevotella* species, which have been shown to stimulate pro-inflammatory cytokines production in several vaginal epithelial cell models *in vitro* ^12–16^. In pregnancy, a shift from *Lactobacillus* dominance towards high-diversity vaginal microbiota associates with increased pro-inflammatory cytokine production and increased cervical vascularisation^9,17^, which in women subsequently experiencing preterm birth, is characterised by excessive activation of the complement cascade^18^.

The mechanisms determining innate immune response to vaginal microbiota remain poorly characterised however innate immune receptor-mediated activation of the transcription factor Nuclear Factor – kappa B (NF-κB) in both immune and epithelial cells is a central feature^10^. In addition to regulating gene transcription of pro-inflammatory mediators, NF-κB mediates expression of key tissue remodelling and pro-labour genes critical to human parturition.^19–21^. Accordingly, untimely activation of innate immune receptors upstream of NF-κB has been associated with pregnancy complications^21^. During murine pregnancy, selective activation of TLR2 and TLR4-dependent signalling pathways in gestational tissues triggers preterm birth^20,22–24^. However, a TLR2-induced preterm birth phenotype is mediated by the co-receptor TLR1, and not TLR6, highlighting the complexity and specificity of innate immune recognition events in the reproductive tract^25^. Despite expressing the TLR2 ligands lipoteichoic acid and teichoic acids in their cell wall, commensal *Lactobacillus* spp. avoid activating pro-inflammatory pathways in the cervicovaginal niche^26^. In some species of lactobacilli S-layer proteins attached to cell wall carbohydrates by non-covalent interactions creating a lattice-like structure across the cell surface can modulate immune responses, including through interactions with anti-inflammatory receptors^27^. In the reproductive tract, these receptors including Siglec 10, which promotes sperm tolerance^28^ and DC-SIGN (CD209), a receptor for HIV,^29,30^ play key roles determining host reproductive fitness. Interactions between microbiota and anti-inflammatory innate immune receptors in the vagina remains a largely unexplored area.

In this study we leverage a large collection of clinical vaginal bacterial isolates to describe the innate immune profile of major vaginal taxa. Our data indicates that *L. crispatus* interacts selectively with DC-SIGN while *L. iners* and BV-associated-bacteria strongly activate TLR2- and TLR4-dependent pro-inflammatory signalling. We show that the unique immunological properties of *L. crispatus* are mediated by their surface layer proteins (SLP), which mask TLR2 ligands from host recognition and mediate interaction with the anti-inflammatory receptor, DC-SIGN. Detection of SLPs in swabs collected during pregnancy associates with vaginal microbiota composition and may act as immunomodulators within the cervicovaginal niche.

## Results

### 1. TLR2 and TLR4 are differentially activated by vaginal taxa associated with non-optimal health states

To understand the role played by vaginal microbiota in regulating the host immune environment, we investigated the capacity of patient-derived vaginal bacterial isolates to induce NF-κB and AP1 activation in HEK cells expressing human TLR2 (HEK TLR2) and TLR4 (HEK TLR4). Both the bacteria and bacterial culture supernatants were tested to evaluate direct activation and activation by secreted immunomodulatory compounds. Neither bacteria nor bacterial supernatants induced activation of the parental HEK-Null cells (Extended Data Fig. 1A, B). Isolates of *L. crispatus*, *L. jensenii* and *L. johnsonii* did not activate the TLR2 reporter cell line, except for *L. jensenii* 19M1 and *L. crispatus* 14M4 and 19N1 which induced low levels of activation (20-40% that of the positive control, Pam2CSK4) (Fig 1A). Two vaginal isolates of *L. gasseri*, 21M4 and 19N2, significantly activated the TLR2 reporter cell line, inducing 56% and 29% activation of the positive control ligand, respectively, while the vaginal isolate, 18M1, and the commercial strain, DSM20077, did not activate TLR2 highlighting strain variability (Fig 1A). Similarly, *L. vaginalis* 10N3, isolated from vaginal swabs, activated the TLR2 reporter cell line whereas the commercially sourced *L. vaginalis* DSM5837, did not (Fig 1A). Culture supernatants containing secreted bacterial compounds of all isolates of *L. crispatus*, *L. gasseri*, *L. jensenii*, *L. johnsonii* and *L. vaginalis* did not activate the TLR2 reporter cell line with the exception of *L. vaginalis* 10N3 (Extended Data Fig. 1C). In contrast to these findings, clinical and commercial *L. iners* isolates and culture supernatants induced TLR2 activation to levels comparable to Pam2CSK4 (Fig 1A and S1C). Similarly, all *G. vaginalis* isolates and culture supernatants activated the TLR2 reporter cell line (Fig 1A and Extended Data Fig. 1C).

**Fig. 1:**
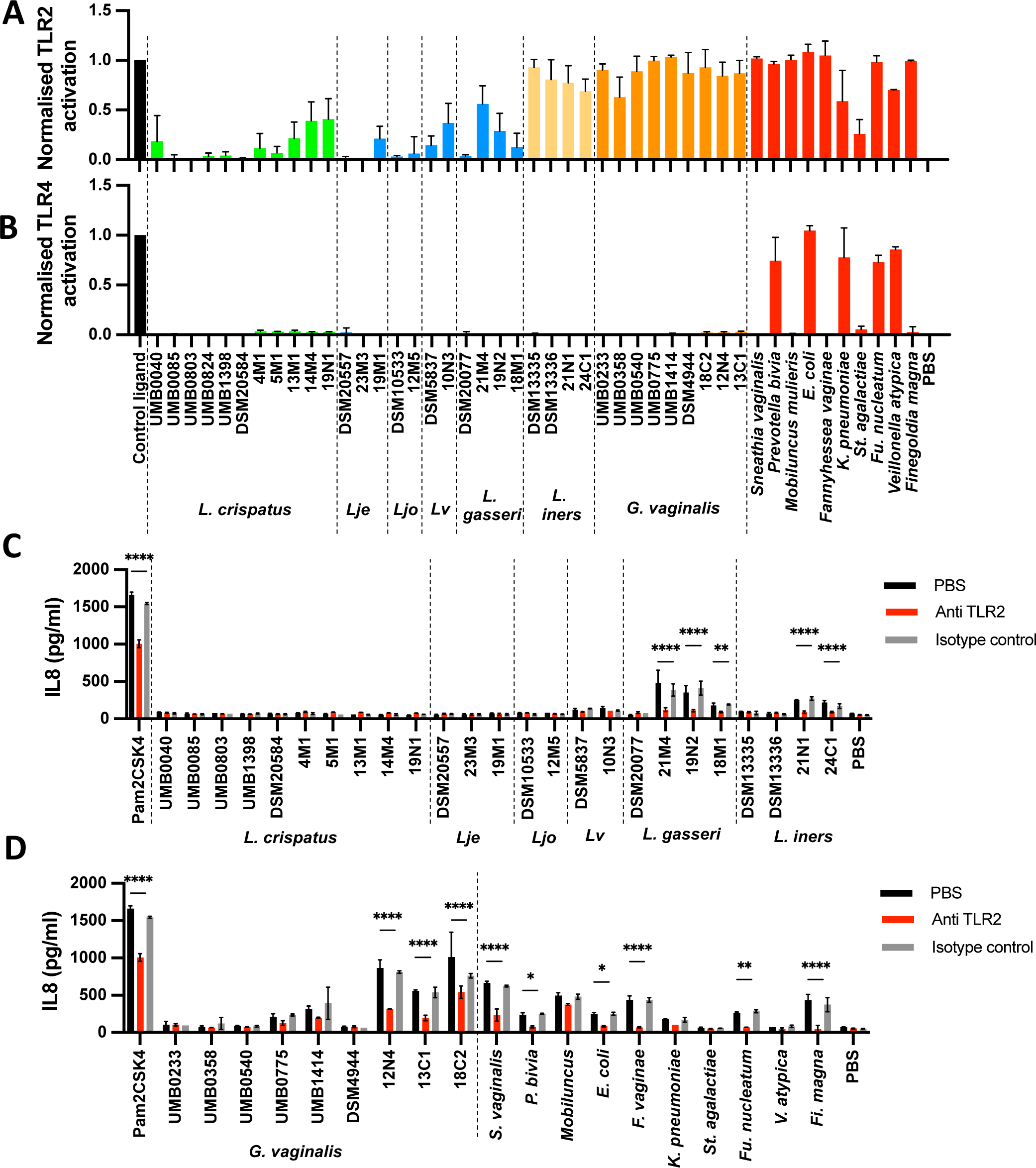
Bacteria associated with preterm birth activate pro-inflammatory signaling pathways R2 and TLR4. HEK TLR2 (A), HEK TLR4 (B) and VK2 vaginal epithelial cells (C, D) were ed with bacteria (MOI=10) overnight. TLR2-dependence of IL-8 production on VK2 cells was ated by pre-incubating the cells for 30min with 1ug/ml anti-TLR2 or isotype control es before stimulation. Pam2CSK4 and LPS EK were used as control ligands for HEK TLR2 EK TLR4 respectively. Lje: Lactobacillus jensenii, Ljo: Lactobacillus johnsonii, Lv: actobacillus vaginalis. Data are mean +/-SD (n=3). * P<0.05, ** P<0.01, *** P<0.001

Isolates of species associated with bacterial vaginosis and vaginal infections including *Sneathia vaginalis, Prevotella bivia, Mobiluncus mulieris, Escherichia coli, Fannyhessea (Atopobium) vaginae, Fusobacterium nucleatum, Klebsiella pneumoniae, Veillonella atypica* and *Finegoldia magna*^5,9,31,32^ strongly activated the TLR2 reporter cell line to levels similar to Pam2CSK4 (Fig 1A and Extended Data Fig. 1C). *Streptococcus agalactiae* induced comparatively lower TLR2 activation (Fig 1A). The only bacteria found to activate the TLR4 reporter cell line were *Prevotella bivia*, *E. coli*, *K. pneumoniae*, *Fu. nucleatum* and *V. atypica* (Fig 1B and Extended Data Fig.1D).

TLR2-dependent immune activation by vaginal taxa was further investigated by examining IL-8 production in VK2 vaginal epithelial cells. While *L. crispatus*, *L. jensenii*, *L. johnsonii* and *L. vaginalis* isolates did not induce IL-8 production in VK2 cells, several isolates of *L. gasseri* and *L. iners* significantly induced TLR2-dependent IL-8 production (Fig 1C). *G. vaginalis* induced IL-8 in a strain-dependent manner, whereas several BV-associated species including *S. vaginalis*, *P. bivia*, *E. coli*, *F. vaginae*, *Fu. nucleatum* and *Fi. magna* were also found to drive IL-8 production via TLR-2 (Fig 1D).

### 2. Vaginal taxa associated with sub-optimal health states activate TLR1/TLR2 pro-inflammatory signalling pathways

TLR2 forms heterodimers with either TLR1 or TLR6 to recognize triacylated and diacylated ligands respectively, and induce pro-inflammatory signalling^33,34^. To investigate TLR2 co-receptor dependency, we evaluated the ability of vaginal bacterial isolates previously shown to stimulate pro-inflammatory pathways, to activate the HEK TLR2 and VK2 cell lines in the presence of anti-TLR1 and TLR6 blocking antibodies. TLR2 signalling induced by isolates of *L. iners* (21N1 and 24C1) and *L. gasseri* (18M1, 19N2 and 21M4) was found to be strictly TLR6 dependent (Fig 2A and 2B). Substantial isolate-to-isolate variability was observed among *G. vaginalis* vaginal isolates. While *G. vaginalis* 12N4 and 13C1 induced both TLR1-and TLR6-dependent TLR2 signalling, *G. vaginalis* 18C2 activated only TLR2/TLR6 signalling (Fig 2A and 2B). Similarly, *S. vaginalis*, *M. mulieris*, *Fi. magna* and *V. atypica* activated only TLR6-dependent TLR2 signalling (Fig 2A). *F. vaginae* and *Fu. nucleatum* were both found to activate TLR1- and TLR6-dependent signalling in HEK TLR2 and VK2 cell lines whereas *P. bivia* activated selectively TLR1-dependent signalling (Fig 2A and 2B).

**Fig. 2:**
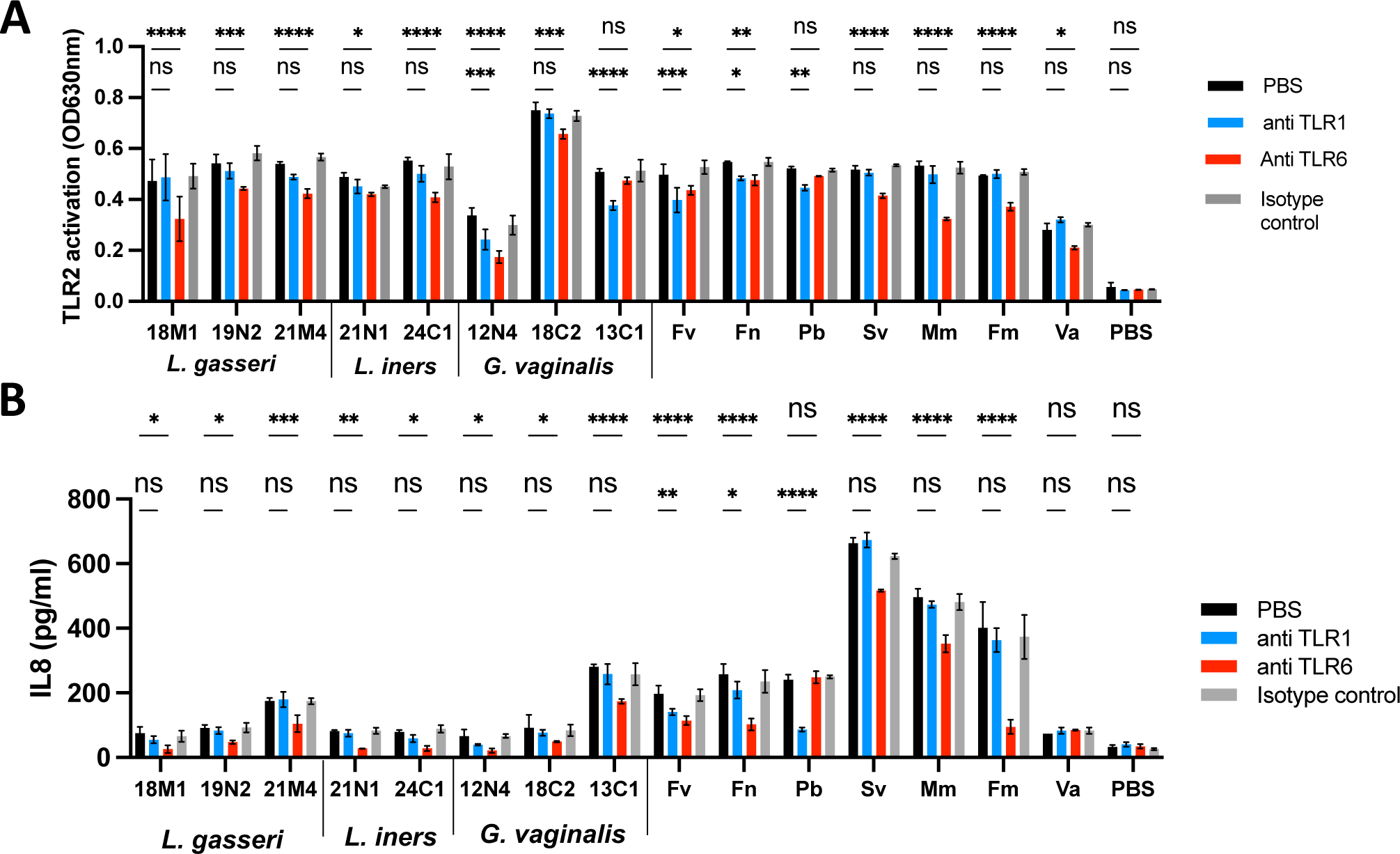
TLR2 co-receptors utilisation by vaginal bacteria. HEK TLR2 (A) and VK2 vaginal al cells (B) were incubated with bacteria (MOI=10) overnight. TLR1 and TLR6-dependence estigated by pre-incubating the cells for 30min with 1ug/ml anti-TLR1, anti-TLR6 or mIgG1 control antibodies before overnight stimulation with the bacteria at MOI=10. Fv: *essea* vaginae, Fn: *Fusobacterium nucleatum*, Pb: *Prevotella bivia*, Sv: *Sneathia vaginalis*, *obiluncum mulieris*, Fm: *Fingoldia magna*, Va: *Veillonella atypica*. Data are mean +/-SD p<0.05, ** p<0.01, *** p<0.005, **** p<0,001

### 3. L. crispatus selectively interacts with the anti-inflammatory receptor DC-SIGN

Anti-inflammatory innate immune receptors are known to play an important role in immune tolerance and immune evasion, including in the context of reproductive biology. We therefore examined the capacity of vaginal taxa to interact with three key anti-inflammatory receptors expressed in the female reproductive tract: Siglec 9^28^, Siglec 10^28^ and Dendritic Cell-Specific Intercellular adhesion molecule-3-Grabbing Non-integrin (DC-SIGN, CD209)^35,36^. Binding assays indicated that none of the lactobacilli interacted with Siglec 9 or Siglec 10 (Fig 3A and 3B). In contrast, apart from *P. bivia*, all BV-associated bacteria displayed binding with Siglec 9 (Fig 3A). *G. vaginalis* binding to Siglec 10 was isolate-dependent with binding observed for *G. vaginalis* UMB0540, UMB1414, 18C2 and to a lesser extent 12N4. No binding was seen between Siglec 10 and *G. vaginalis* UMB0233, UMB0358, UMB0775 and DSM4944 (Fig 3B). Interactions were also observed between Siglec10 and *S. vaginalis*, *E. coli*, *F. vaginae* and *St. agalactiae* (Fig 3B).

**Fig. 3:.**
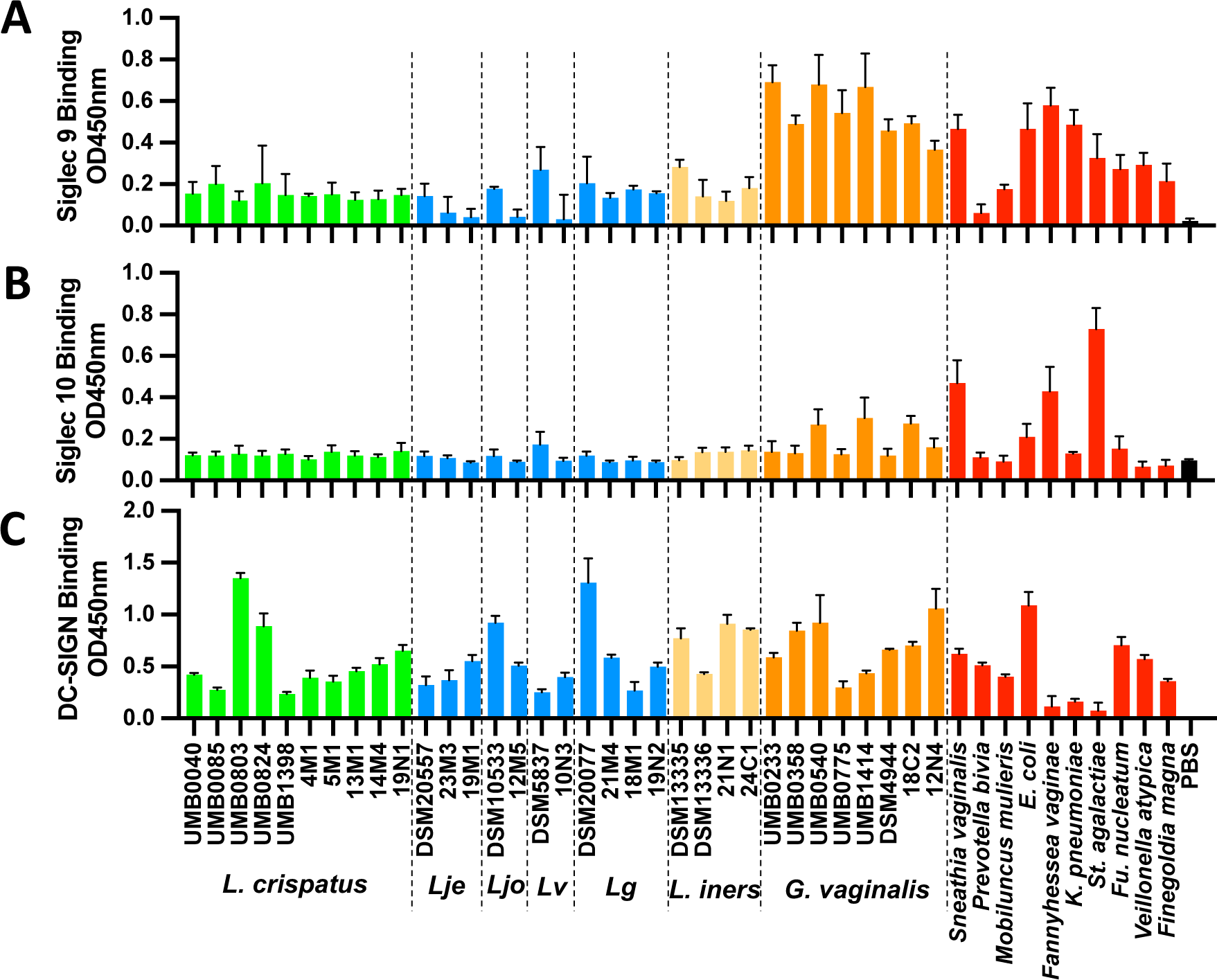
Differential binding to inhibitory lectins by bacteria associated with term d preterm birth. Bacteria (106 per well) were coated in 96-well plates and tested for ir capacity to bind Siglec 9-Fc (A), Siglec 10-Fc (B) and DC-SIGN-Fc (C). Lje: L. senii, Ljo: L. johnsonii, Lv: Limosilactobacillus vaginalis, Lg: L. gasseri. Data are an +/-SD (n=3).

Binding of vaginal taxa to DC-SIGN was highly variable between isolates. *L. crispatus* UMB0803 and UMB0824, *L. johnsonii* and *L. gasseri* DSM20077 binding to DC-SIGN was relatively high compared to other isolates (Fig 3C). *G. vaginalis* isolates also interacted with DC-SIGN, however, no binding was observed for *F. vaginae*, *K. pneumoniae* and *St. agalactiae.* DC-SIGN binding was also observed for *S. vaginalis*, *P. bivia*, *M. mulieris*, *E. coli*, *F. nucleatum*, *V. atypica* and *Fi. magna* (Fig 3C). Consistent with previously described DC-SIGN carbohydrate specificity^37–39^, interactions between bacterial isolates and DC-SIGN were inhibited in presence of glucose, mannose and EDTA but not galactose (Extended Data Fig. 2A and 2B).

### 4. S-layer proteins are selectively expressed by L. crispatus and shield TLR2 ligands

The cell wall of lactobacilli is known to contain TLR2 ligands such as lipoteichoic acids and lipoproteins^26^. S-layer proteins (SLPs), expressed by some lactobacilli, form a 2D crystalline array on the cell surface that mediates interactions with other cells including the potential shielding of cell wall constituents from detection^26,27^. Using an adapted lithium chloride-based extraction method^40^, we next examined SLPs expression in *L. crispatus* and *L. iners* isolates. S-layer proteins within the expected relative molecular weight range could be isolated from all *L. crispatus* isolates^27^ (Fig 4A, Extended Data Fig.3A and 3B). *L. crispatus* UMB0040, 4M1, 19N1 and 14M4 were found to predominately express the 60kDa SLP1 (Fig 4A) while *L. crispatus* 5M1, 13M1, UMB0085, UMB803, UMB824 and UMB1398 expressed the 45kDa SLP2 (CsbA) (Fig 4A, Extended Data Fig. 3A and 3B). Aggregation Promoting Factors (Afp), which share sequence homology with SLPs, have also been isolated from *L. johnsonii* and *L. gasseri* using lithium chloride extraction^41,42^. Proteins of corresponding molecular weights were detected in *L. johnsonii* DSM10533 (42kDa) and *L. gasseri* DSM20077 (32kDa) (Extended Data Fig. 3B). However, SLPs or Afp were not detected in any other *L. johnsonii*, *L. jensenii*, *L. gasseri* or *L. iners* vaginal isolates. (Fig 4A and Extended Data Fig. 3A).

**Fig. 4:**
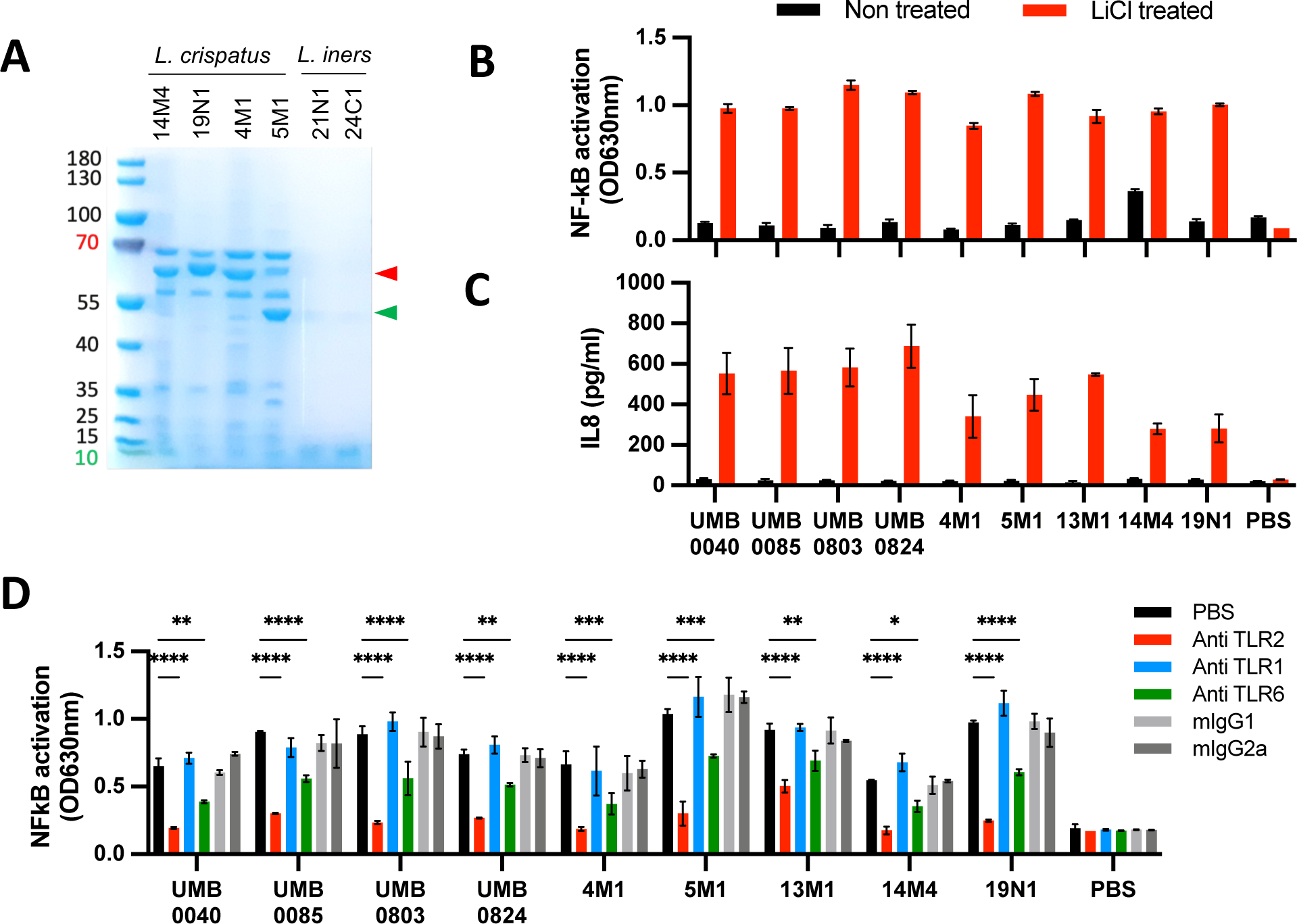
SLPs mask TLR2 ligands of *L. crispatus* and prevent TLR2-dependent pro-matory pathways activation. (A) SDS-PAGE gel of crude SLPs extracted with M from *L. crispatus* and *L. iners* isolates. 10ug total proteins were loaded per lane. EK-TLR2 cells were stimulated for 16 h with bacteria (MOI=10) and NF-κB ion was determined by measuring alkaline phosphatase activity and reading O.D. nm. (C) VK2 vaginal epithelial cells were stimulated for 16 h and IL-8 release in pernatant was quantified by ELISA. (D) HEK TLR2 cells were stimulated with chloride-treated bacteria (MOI=10) for 16 h and IL-8 release in the supernatant uantified by ELISA. TLR1, TLR2 and TLR6 dependence was investigated by pre-ting cells for 30 min at 37 °C with 1 μg/ml of anti-TLR1, anti-TLR2, anti TLR6 or and mIgG2b isotype control antibodies. (B, C, D) A representative figure of three ndent experiments is shown (mean +/-SD). * p<0.05, ** p<0.01, *** p<0.005, ****

As previously reported, *L. crispatus* isolates did not activate the HEK TLR2 reporter cell line. However, removal of SLPs from the cell surface using LiCl led to strong activation of the HEK TLR2 reporter cell line (Fig 4B) and induction of IL-8 production by VK2 vaginal epithelial cells (Fig 4C). Furthermore, activation of the TLR2 reporter cell line was shown to be TLR2- and TLR6-dependent, but not TLR1-dependent, for all *L. crispatus* isolates tested (Fig 4D). SLPs purified from *L. crispatus* isolates did not activate the TLR2 reporter cell line (Extended Data Fig. 3C).

### 5. SLPs from L. crispatus are ligands of DC-SIGN

We next investigated the capacity of SLPs isolated from different vaginal taxa to interact with the anti-inflammatory receptor, DC-SIGN. SLP1 purified from *L. crispatus* UMB0040, 4M1, 14M4 and 19N1 and SLP2 from *L. crispatus* DSM20584, UMB0824, UMB1398, 5M1 and 13M1 were shown to bind DC-SIGN by western blot (Fig 5A and Extended Data Fig. 4A). Similarly, SLP isolated from *L. johnsonii* was also capable of binding DC-SIGN. Binding of DC-SIGN to *L. crispatus* SLPs was inhibited in the presence of glucose, mannose and EDTA but not galactose for all the strains tested (Fig 5B and Extended Data Fig. 4C), indicating glycan-mediation of SLPs-DC-SIGN interactions. In contrast, lithium chloride-extracts from *L. iners* DSM13335 and DSM13336, *L. gasseri* DSM20077, *L. jensenii* DSM20557 and *L. vaginalis* DSM5487 did not bind DC-SIGN (Extended Data Fig. 4A and 4B).

**Fig. 5:**
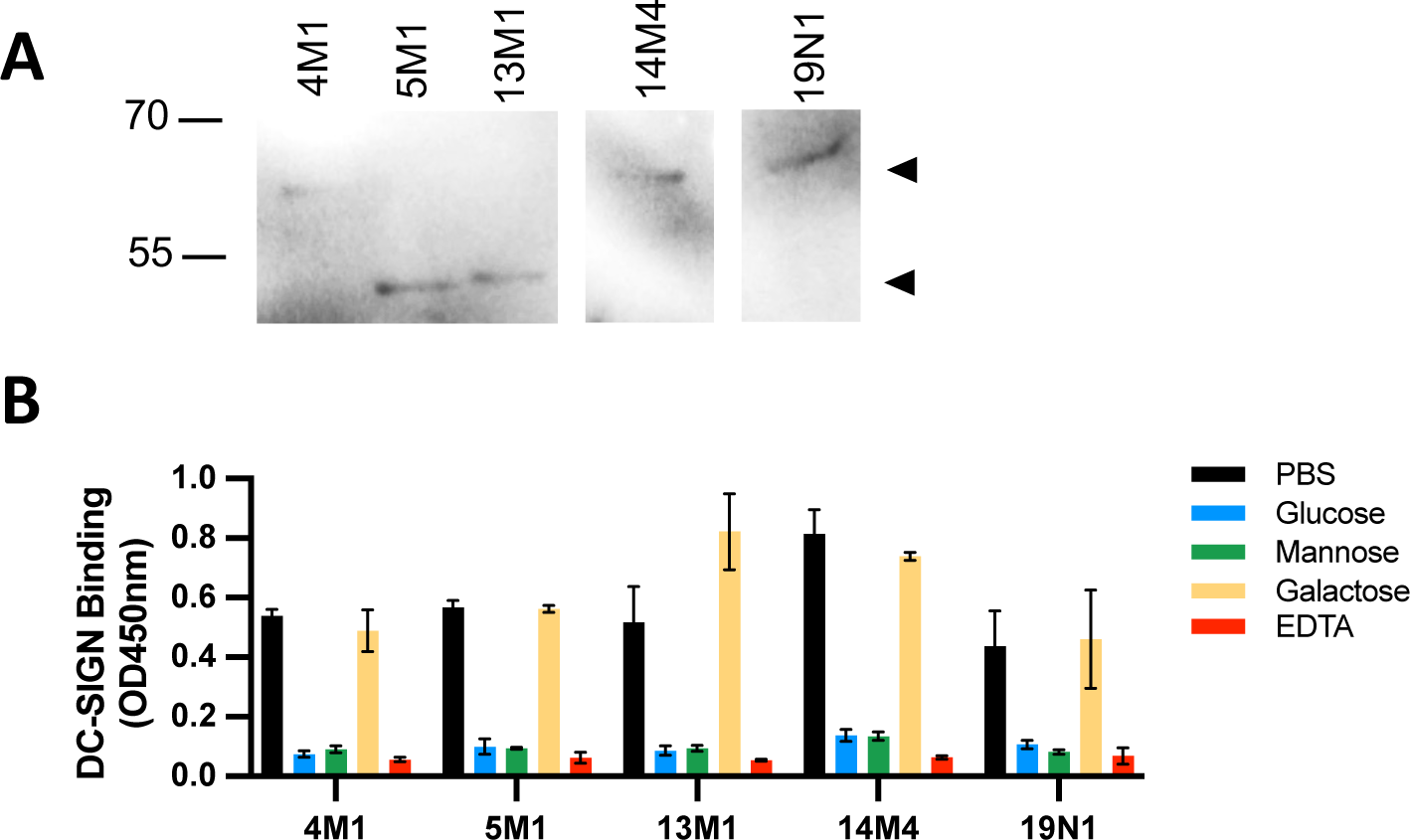
SLPs are ligands of DC-SIGN. (A) Western blotting results for DC-SIGN-Fc ing to purified SLPs isolated from vaginal L. crispatus strains (one representative of e replicates), (B) Purified SLPs isolated from vaginal L. crispatus strains (1ug per well) coated in 96-well plates and tested for their capacity to bind DC-SIGN-Fc. DC-SIGN-1 μg/ml) was pre-incubated or not with 20 mM EDTA or 40 mM glucose, mannose and ctose and allowed to react with the SLPs for 2 h at RT. Bound proteins were detected g a biotin-conjugated anti-IgG Fc specific antibody and avidin-HRP and reading O.D. 50 nm. Data are mean +/-SD (n=3).

### 6. Detection of SLPs in human vaginal secretions and association with microbiota composition

Surface layer proteins from other bacterial species have been reported to be shed freely into the environment^43,44^ or released via incorporation in extracellular vesicles^45^. We therefore investigated if SLPs could be detected in cervicovaginal fluids samples collected from a cohort of pregnant women with microbiota compositions dominated by *L. crispatus* (CST I, n=19) or *L. iners* (CST III, n=10) or *Lactobacillus-*depleted, high diversity compositions enriched for BV-associated bacteria (n=10). SLPs were detected in 15 out of 19 CST I samples (78%), but none of the CST III samples examined (Fig 6A and 6B). Two out of 10 CST IV samples (20%) were positive for SLPs, of which both samples contained *L. gasseri* as a minor community member (Fig 6B and Extended Data Fig. 5). DC-SIGN binding was demonstrated for all SLPs-positive cervicovaginal fluids samples apart from three (B007, B090 and BG003) (Fig 6B and 6C).

**Fig. 6:.**
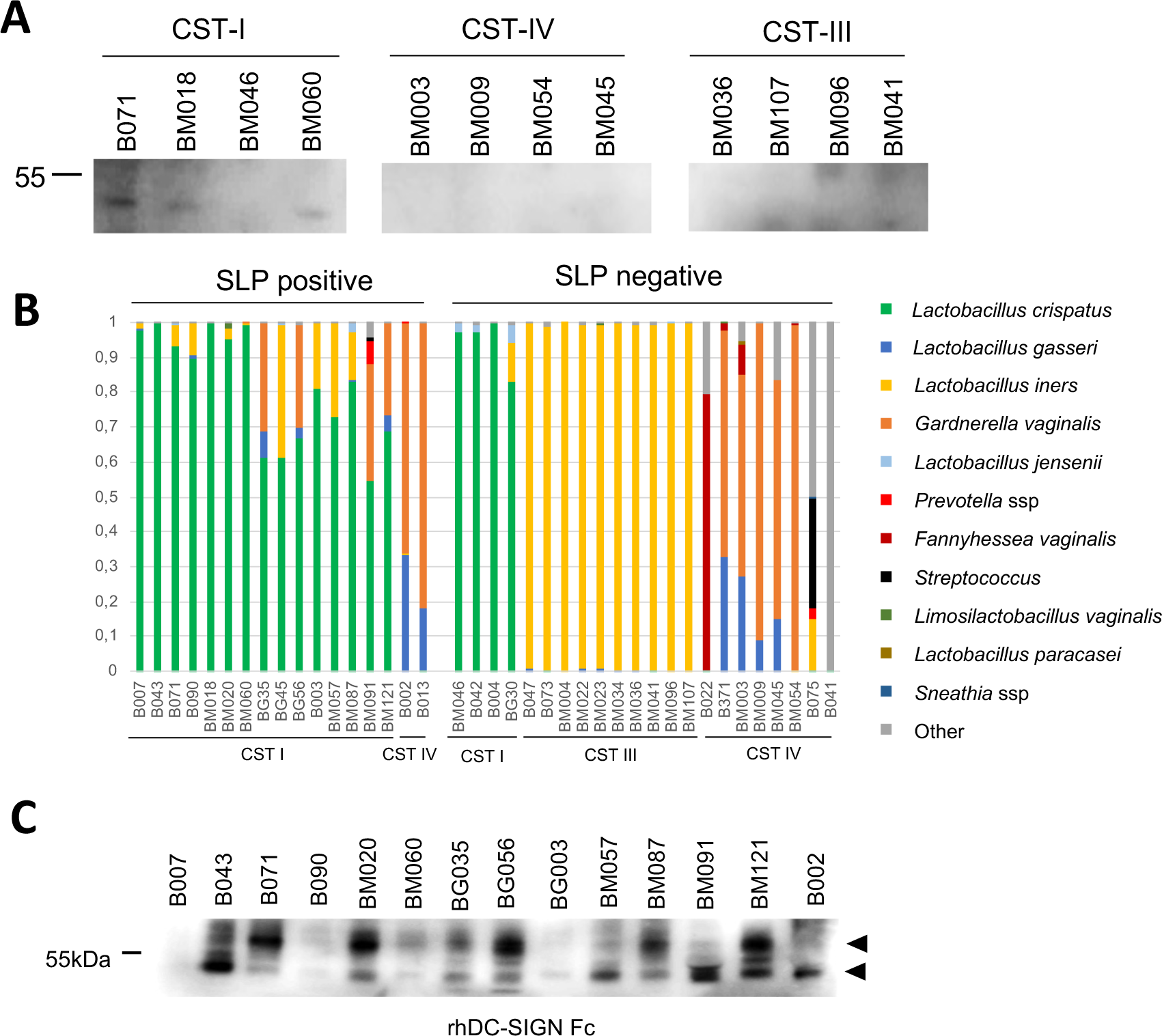
SLPs are detected in cervico-vaginal fluids. (A) Western blotting results anti-Surface Layer Protein antibody binding to purified cervico-vaginal fluid ples, (B) Bacterial composition of the cervico-vaginal fluid samples determined 16S rRNA gene sequencing (C) Western blotting results for DC-SIGN-Fc ding to CVF samples.

## Discussion

A loss of *Lactobacillus* species from the vaginal microbiota and a concurrent increase in bacterial diversity is associated with inflammation, predisposition to acquiring STIs and UTIs and during pregnancy, an increased risk of preterm birth^8–10,46^. However, host inflammatory response to vaginal taxa is highly heterogenous yet closely linked to pathophysiology^18,47,48^. For example, we have recently shown that during pregnancy women experiencing preterm birth are more likely to be carrying high diversity vaginal bacterial community compositions that are often, but not always, accompanied by high innate immune response^18^. By characterising and comparing innate immune signalling of major commensal and pathogenic bacteria isolated from human vaginal samples, the current study provides important insight into how vaginal bacteria differentially stimulate host-cell inflammatory response. Moreover, we report a mechanism by which *L. crispatus*, a commensal widely associated with optimal vaginal health, avoids stimulation of innate immune response via SLPs-shielding of TLR2/4 ligands and selective interacts with the anti-inflammatory receptor DC-SIGN.

Using cell reporter systems, we found that clinical isolates representative of prevalent vaginal commensal and pathogenic species induce TLR signalling in a highly heterogenous manner. While *Lactobacillus* species, particularly those associated with optimal vaginal and reproductive health such as *L. crispatus* did not activate TLR2 or TLR4 pathways, species associated with sub-optimal vaginal health predominately activated TLR2 signalling in an isolate-dependent manner. This included *L. iners*, which often acts as a transitional coloniser of the cervicovaginal niche^49^ and *G. vaginalis*, a species linked to BV and increased risk of preterm birth^9^. This variability appears to extend to vaginal epithelial cells where differential production of IL-8 was observed in response to exposure to differing vaginal taxa. These findings provide insight into recent *in vivo* studies which have observed high patient-to-patient variability in inflammatory response to vaginal species^9,18^. These studies have been limited to the characterisation of vaginal microbiota composition to species taxonomy. Our data indicates that different strains of vaginal commensals and pathogens interact with TLR and anti-inflammatory receptors to differing degrees, which would partly account for the observed heterogeneity in host inflammatory response.

Isolate-to-isolate variability for *G. vaginalis* was also observed for binding of anti-inflammatory receptors, Siglec 9 and Siglec 10. Subspecies of *G. vaginalis* have been shown to differ in terms of virulence factors expressed, biofilm formation capacity and/or the presence of a polysaccharide capsule^25,42^. Such structural variations could underlie differing immune profiles as observed in this study, with exposure of anti-inflammatory molecular motifs acting as ligands for Siglec 9 and Siglec 10 thus modifying tolerance of potentially pathogenic species. Consistent with these findings, specific sub-species or strains of *G. vaginalis* have recently been implicated in increased risk of preterm birth^50,51^. We also observed variability of vaginal isolate binding to the anti-inflammatory receptor DC-SIGN. Despite this, strongest binding was observed for isolates of *L. crispatus*, the binding of which was shown to be mediated by specific glycan interactions. These findings are in agreement with variability observed in the predicted carbohydrate binding protein repertoire of vaginal pathogen and commensal species^52^ and highlight the importance of examining the relationship between strain-level resolution of the vaginal microbiota, host inflammatory response and clinical outcomes.

Our findings also highlight SLPs as important mediators of host cell inflammatory response to *L. crispatus*. By shielding ligands contained within the cell wall of *L. crispatus*, SLPs were shown to prevent activation of pro-inflammatory signalling via TLR2 and moreover to be ligands for DC-SIGN. In our assays SLP1 and SLP2 seemed to play a redundant role in preventing TLR2 activation, however SLP2 has previously been shown to contain an amino-terminal domain that binds to type I and IV collagen and a carboxyl-terminal domains that modulates cell wall binding^53,54^. SLPs have been shown to dynamically regulate immune and inflammatory response to bacteria in other species via the modulation of bacterial cell wall permeability^55^. Our observation that SLPs are found not only on the surface of *L. crispatus* but also released in CVF indicates that secreted SLPs may contribute to the anti-inflammatory environment of the vaginal niche when dominated by *L. crispatus*.

In summary, our findings indicate that the vaginal lactobacilli associated with optimal health interact selectively with a very restricted subset of anti-inflammatory receptors through their Surface Layer Proteins, both *in vitro* and in cervico-vaginal fluids. We propose that these interactions shape and stabilize the maternal immune environment, preventing preterm parturition and other adverse health outcomes. Furthermore, bacterial strains activating TLR2 and TLR4 but also Siglec 9 and Siglec 10 are associated with bacterial vaginosis and adverse pregnancy outcome, indicating that some anti-inflammatory receptors may promote immune evasion and co-colonization by pathogenic strains rather than play a protective role. Overall, our data provide a rational ground to guide the selection and development of vaginal live biotherapeutic products with optimal innate immune properties, as well as host-targeted immunomodulatory therapeutics for the prevention and treatment of adverse women’s health outcomes.

## Material and methods

### Patient recruitment and sample collection

Pregnant women at risk of preterm birth were recruited from preterm birth prevention clinics from Queen Charlotte’s and Chelsea Hospital (QCCH) between June 2018 and June 2022. The study was performed under the Ethics approval REC 14/LO/0328 as part of the Vaginal Microbiome and Metabolome in Pregnancy (VMET 2) Research Study and approved by the NHS Health Research Authority (London -Stanmore Research Ethics Committee). All patients provided written informed consent to donate specimens.

Cervicovaginal Fluid (CVF) was first sampled using a BBL™ Culture Swab™ MaxV Liquid Amies swab (Becton, Dickinson and Company, Oxford, UK). Swabs were placed on ice immediately and stored at –80°C until use. A soft cup (flexTM, menstrual disc) was then placed past the vaginal canal for 20 minutes, retrieved and the weight of the CVF collected and recorded. CVF from soft cups was mixed with 5ml PBS per gram CVF and collected. Diluted CVFs were centrifuged at 4°C for 10min at 16,000 x *g* before the supernatant was collected and stored at -80°C.

### Bacterial strains and culture conditions

Patient derived bacterial isolates used in this study are described in Extended Data table 1. Briefly, bacteria were cultured in their corresponding medium overnight at 37°C. All the bacteria (except *E. coli* and *K. pneumoniae*) were grown anaerobically (10% CO_2_, 10% hydrogen, 80% nitrogen; 70% humidity) and were washed in degassed HBSS. *E. coli* and *K. pneumoniae* were grown aerobically and washed with PBS. Aerobic liquid cultures were grown in a shaking incubator at 200 rpm. The bacteria were quantified as previously described^56^. The bacterial pellet was stored at -20°C until further use.

### S-layer protein purification

Extraction of S-layer proteins was performed as previously described^40^. Briefly, bacterial pellets of 50ml overnight cultures were washed with PBS and resuspended in LiCl 5M at 4°C for 15 min under stirring. The supernatants were harvested by centrifugating 10min at 3000 x *g* and dialyzed against water overnight at 4°C in Snakeskin dialysis tubing cut off 10kDa (68035, ThermoFisher Scientific). Precipitated S-layer proteins were recovered by centrifugation at 20,000 *x g* for 20min. The proteins were solubilized in LiCl 5M and further purified by size exclusion chromatography using Sephacryl S200R (S200HR-250ML, Sigma Aldrich). The S-layer proteins were characterized by SDS-PAGE and stained with Coomassie Blue. A single band was obtained for all the strains.

### Binding of Fc-tagged lectins

Bacteria (10^6^/well in isopropanol) or purified S-layer proteins (1 μg/well in isopropanol) were coated on 96-wells high binding plates (Sarstedt). DC-SIGN-Fc (10200-H01H-SIB-100ug, Sino Biological), Siglec-9-Fc (1139-SL-050, BioTechne) or Siglec-10-Fc (2130-SL-050, BioTechne) fusion proteins (1 μg/ml in PBS, 1 mM CaCl_2_, 1% w/v BSA) were precomplexed with anti-human Fc specific antibodies-HRP (A18817, ThermoFisher Scientific, 2:1 ratio lectin:antibody) and pre-incubated with 20 mM EDTA or 40 mM glucose, mannose or galactose (Tokyo Chemical Industry UK Ltd) when indicated. The recombinant lectins were allowed to bind to bacterial cells or SLPs for 2h at RT (in 50 μl). Wells were washed once with PBS and the plates were incubated with a TMB (ES001-500ML, EMD). The reaction was stopped with HCl 1M and the plates were read at 450nm.

### TLR2 and TLR4 reporter cell lines experiments

The HEK-Blue hTLR2 and HEK-Blue hTLR4 (InvivoGen), derivatives of HEK293 cells that stably express the human TLR2 and TLR4 genes respectively, along with a NF-κB-inducible reporter system (secreted alkaline phosphatase) were maintained in Dulbecco’s modified Eagle’s medium (DMEM, Sigma Aldrich) containing 10% v/v Fetal Bovine Serum (FBS, Sigma Aldrich), 2 mM L-glutamine, 100 U/ml penicillin and 100 μg/ml streptomycin (Sigma). HEK Null1 cells (InvivoGen) were used as parental cell line. Reporter cells (5 × 10^4^/well) were stimulated with bacteria (MOI=10) or purified SLPs (from 10 μg/ml to 80ng/ml) for 16 h, after which alkaline phosphatase activity was measured by mixing 10 μl of the culture supernatant and 90 μl of Quanti-Blue (InvivoGen) and reading O.D. at 630 nm.

### VK2 vaginal epithelial cells experiments

The VK2/E6E7 cells (ATCC) were maintained in Keratinocyte Serum Free Medium (KSFM, Gibco) containing 100 U/ml penicillin and 100 μg/ml streptomycin (Sigma). The VK2/E6E7 cells (3 × 10^4^/well) were stimulated with bacteria (MOI=10) or purified SLPs (from 10 μg/ml to 80ng/ml). After 16 h, IL-8 was assayed in the culture supernatant using a commercially available kit (88-8086-77, ThermoFisher Scientific). To investigate TLR1, TLR2 and TLR6 dependence, VK2/E6E7 cells were pre-incubated for 30 min at 37 °C with 1 μg/ml of anti-hTLR1 (mabg-htlr1, InvivoGen), anti-hTLR2 (MAB2616, R&D Systems) and anti-hTLR6 (mabg-htlr6, InvivoGen) antibody or mIgG1 (02-6100, ThermoFisher Scientific) and mIgG2b (14-4732-85, ThermoFisher Scientific) isotype controls.

### Western blot analysis

A total of 1 ug surface layer protein was added to Laemmli 10mM DTT buffer followed by denaturing at 95 °C for 5 min. For soft cup samples, 10ul sample was used per lane. The proteins were separated by SDS–PAGE and transferred onto PVDF membranes. Blots were incubated with anti-Surface Layer Protein antibody (BS3797, Bioss) and DC-SIGN-Fc (10200-H01H-SIB-100ug, Sino Biological) at 1ug/ml in TBS-T 5% BSA. As secondary antibodies, anti-rabbit-IgG-HRP (1:2000) (Invitrogen) or anti-human-IgG-HRP (1:2000) (ThermoFisher Scientific) were used respectively. ECL signal was recorded on the ChemiImager LAS4000 and data were analysed with ChemiImager LAS4000 software (GE Healthcare).

### DNA extraction and 16S rRNA gene sequencing

DNA extraction from the BBL™ CultureSwab™ was performed as previously described^57^. The V1-V2 hypervariable regions of bacterial 16S rRNA genes were amplified using a forward primer set (28f-/YM) mixed to a 4:1:1:1 ratio of the following primers: 28F-Borrellia GAGTTTGATCCTGGCTTAG, 28F-Chlorlex GAATTTGATCTTGGTTCAG, 28F-Bifido GGGTTCGATTCTGGCTCAG, 28F-YM GAGTTTGATCNTGGCTCAG; and a reverse primer that consisted of 388R GCTGCCTCCCGTAGGAGT. Sequencing was performed at Research and Testing Laboratories (RTL Genomics, Texas, USA). Cutadapt (version 2.8) was used for primer sequences trimming^58^. ASV (amplicon sequence variant) counts for each sample were computed using the QIIME2 bioinformatics pipeline (version2022.2.1)^59^. DADA2 (version 2022.2.0)^60^ was utilised for denoising with forward and reverse read truncation legnths at 210 and 175 nucleotides respectively. Finally, the STIRRUPS database^61^ was used for taxonomic classification of ASVs. At species level, samples were classed using the VALENCIA centroid classification algorithm into five community state types^62^.

### Statistical analysis

Prism software (Graphpad Software) was used to perform statistical tests and to generate graphs. Data are presented as mean ± s.e.m. and P values were calculated using two-way ANOVA and Tukey’s post-test.

## Acknowledgements

We would like to thank all women who have participated in this study and members of the Women’s Health Research Centre who facilitated and coordinated study recruitment and sample collection. *L. crispatus* UMB0040, UMB0085, UMB0803, UMB0824 and UMB01398, *G. vaginalis* UMB0233, UMB0358, UMB0540, UMB0775 and UMB1414 and *S. agalactiae* UMB0776 were kindly gifted by the Wolfe Lab. This work was funded by the March of Dimes European Preterm Birth Research Centre at Imperial College London and supported by the National Institute of Health Research (NIHR) Imperial Biomedical Research Centre (BRC). L.S. is supported by The Parasol Foundation Clinical Senior Lecturer scheme, and B.G.M. is supported by The Parasol Foundation Fellowship Scheme and The Rosetrees Trust. A.D. is funded by an Imperial College Research Fellowship.

## Authors contributions

A.D., J.R.M., L.S., P.R.B. and D.A.M. designed research; A.D., I.K., B.G.M., C.C., D.B.R. and Q.Y.T. performed research; L.R. isolated and cultured the bacterial strains; S.N. processed the sequencing data; data processing, analysis, and interpretation was performed by A.D., I.K., L.R., B.G.M., C.C., D.B.R., Q.Y.T., J.R.M., L.S., P.R.B. and D.A.M.; and A.D. and D.A.M. wrote the first draft of the manuscript. All authors critically reviewed, read and approved the final manuscript.

## Competing interests

The authors declare no competing interests.

---

**Extended Data Fig. 1:**
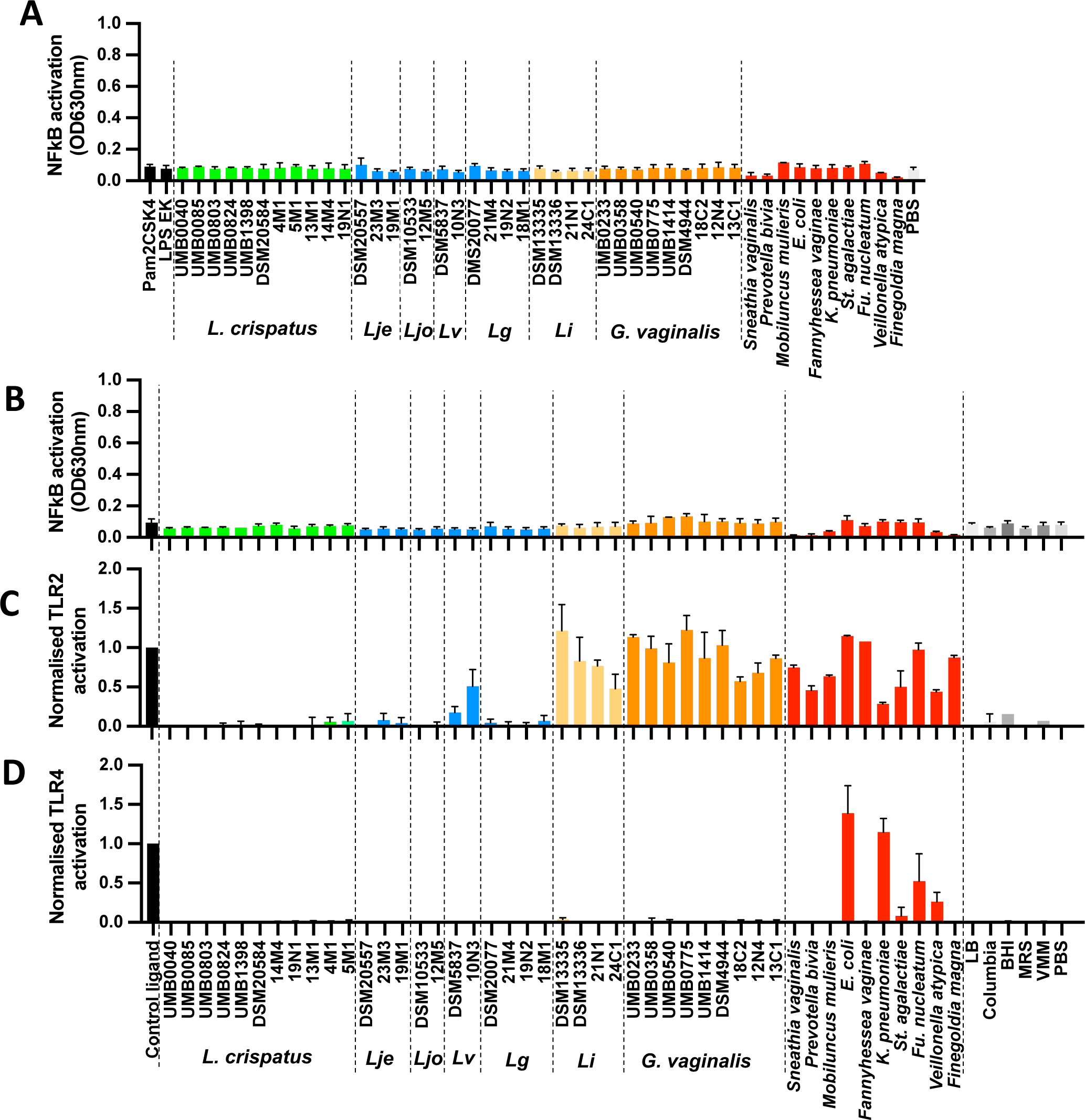
Bacteria and bacterial culture supernatant associated with preterm birth activate pro-inflammatory signaling pathways via TLR2 and TLR4. (A) The HEK Null cells were incubated overnight with the bacteria (MOI=10) and activation of the NFkB and AP1 dependent pro-inflammatory pathways was measured. HEK Null1 parental cell line (B) HEK TLR2 (C) and HEK TLR4 (D) were incubated with 10ul of bacterial culture medium overnight. Pam2CSK4 and LPS EK were used as control ligands for HEK TLR2 and HEK TLR4 respectively. Lje: *Lactobacillus jensenii*, Ljo: *Lactobacillus johnsonii*, Lv: *Limosilactobacillus vaginalis*, *Lg: L. gasseri,* Li: *L. iners*. Data are mean +/-SD (n=3).

**Extended Data Fig. 2:**
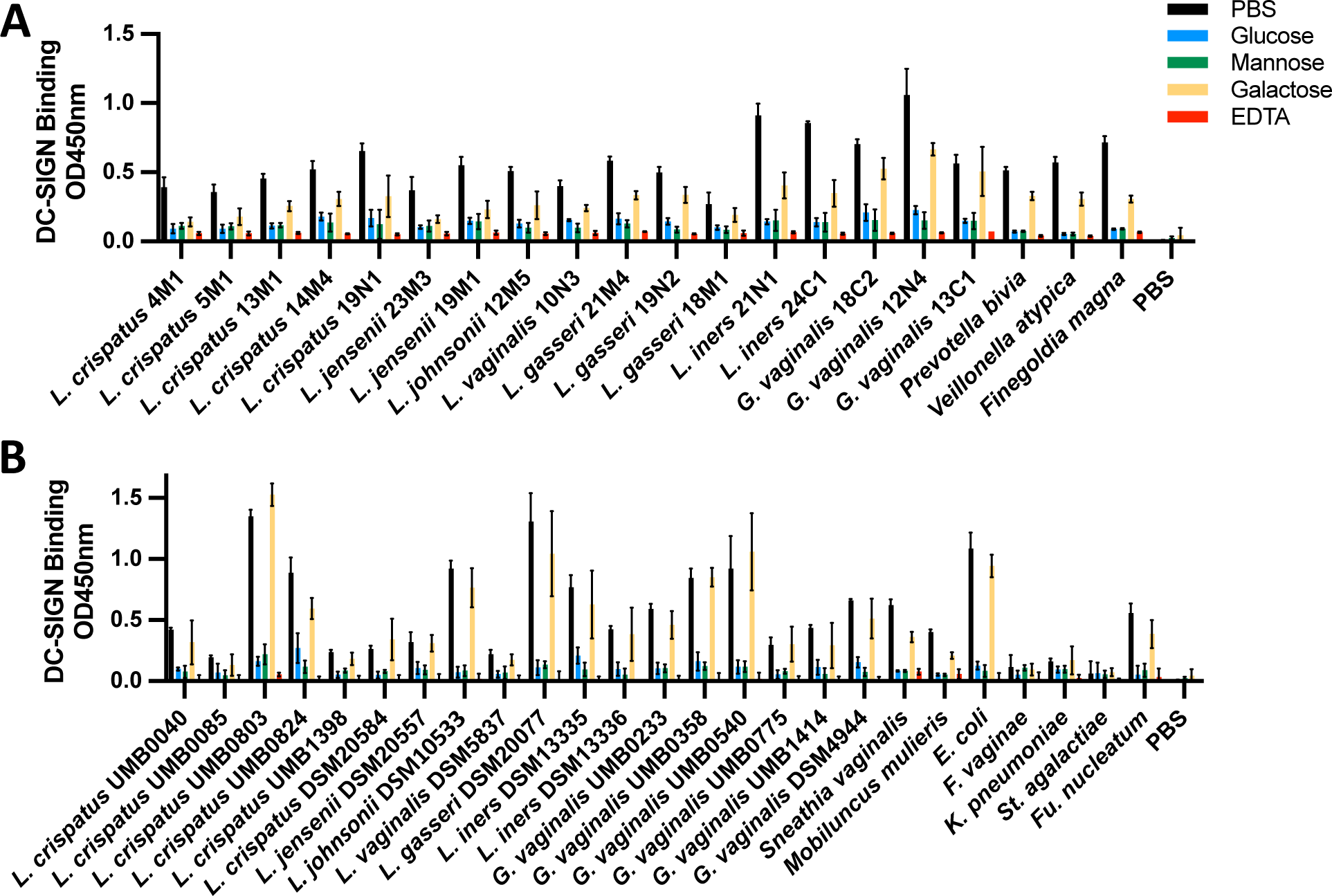
DC-SIGN binding to the bacteria is glycan-dependent. Bacteria ed from vaginal swabs. (A) and commercial strains and bacteria isolated from urinary infection patients (B) (10^6^ bacteria per well) were coated in 96-well plates and tested for capacity to bind DC-SIGN-Fc. DC-SIGN-Fc (1 μg/ml) was pre-incubated or not with M EDTA or 40 mM glucose, mannose and galactose and allowed to react with the ria for 2 h at RT. Bound proteins were detected using a biotin-conjugated anti-IgG Fc fic antibody and avidin-HRP and reading O.D. at 450 nm. Data are mean +/-SD (n=3).

**Extended Data Fig. 3:**
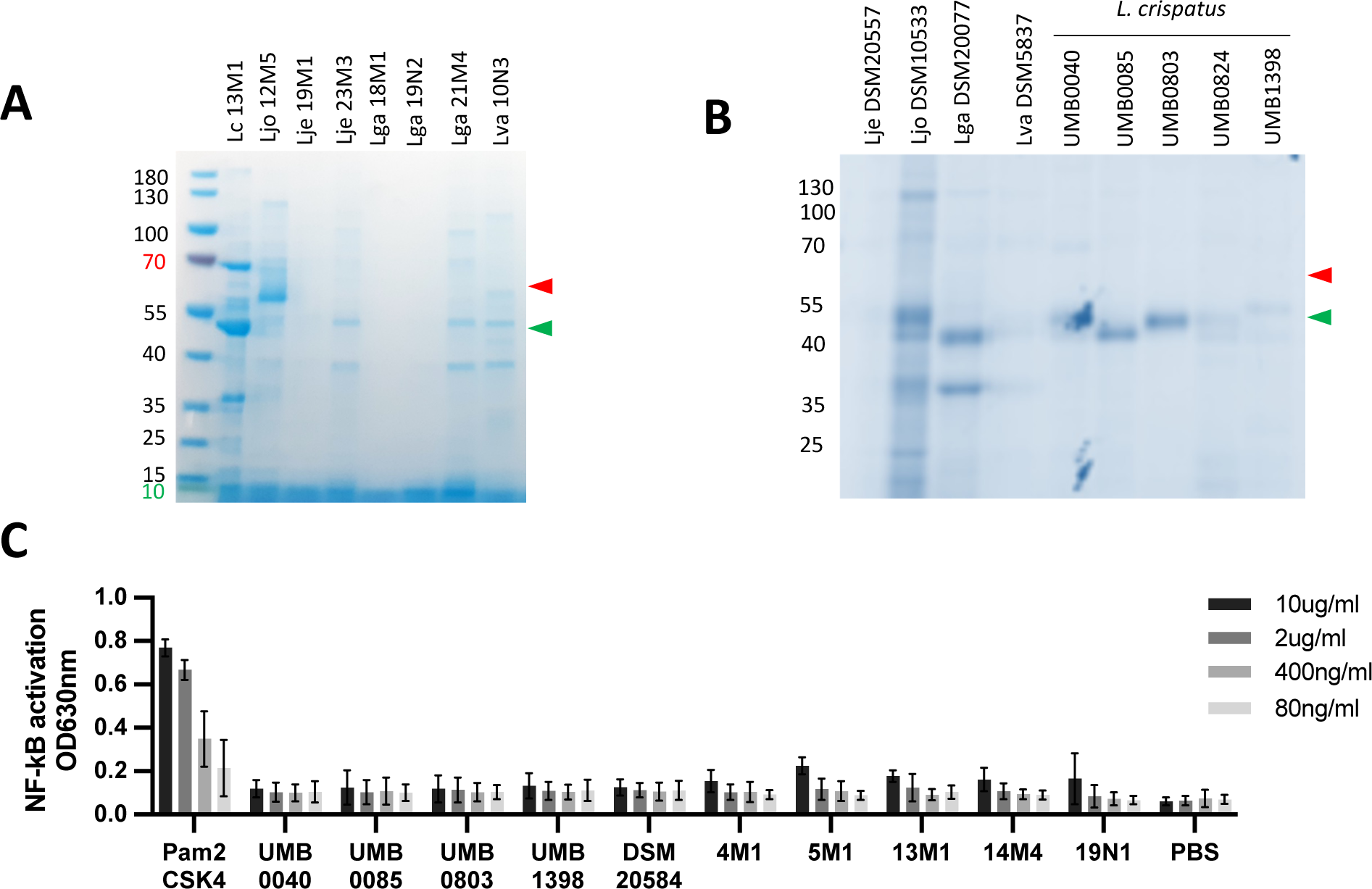
SLPs isolated from *L. crispatus* don’t activate TLR2 signaling. (A) PAGE gel of crude SLPs extracted with LiCl 5M from vaginal lactobacilli isolates. 10ug proteins were loaded per lane. (B) SDS-PAGE gel of crude SLPs extracted with LiCl 5M commercial and urine lactobacilli isolates. 10ug total proteins were loaded per lane. (C) TLR2 cells were stimulated for 16 h with purified SLPs from the *L. crispatus* isolates and B activation was determined by measuring alkaline phosphatase activity and reading at 630 nm. Data are mean +/-SD (n=3).

**Extended Data Fig. 4:**
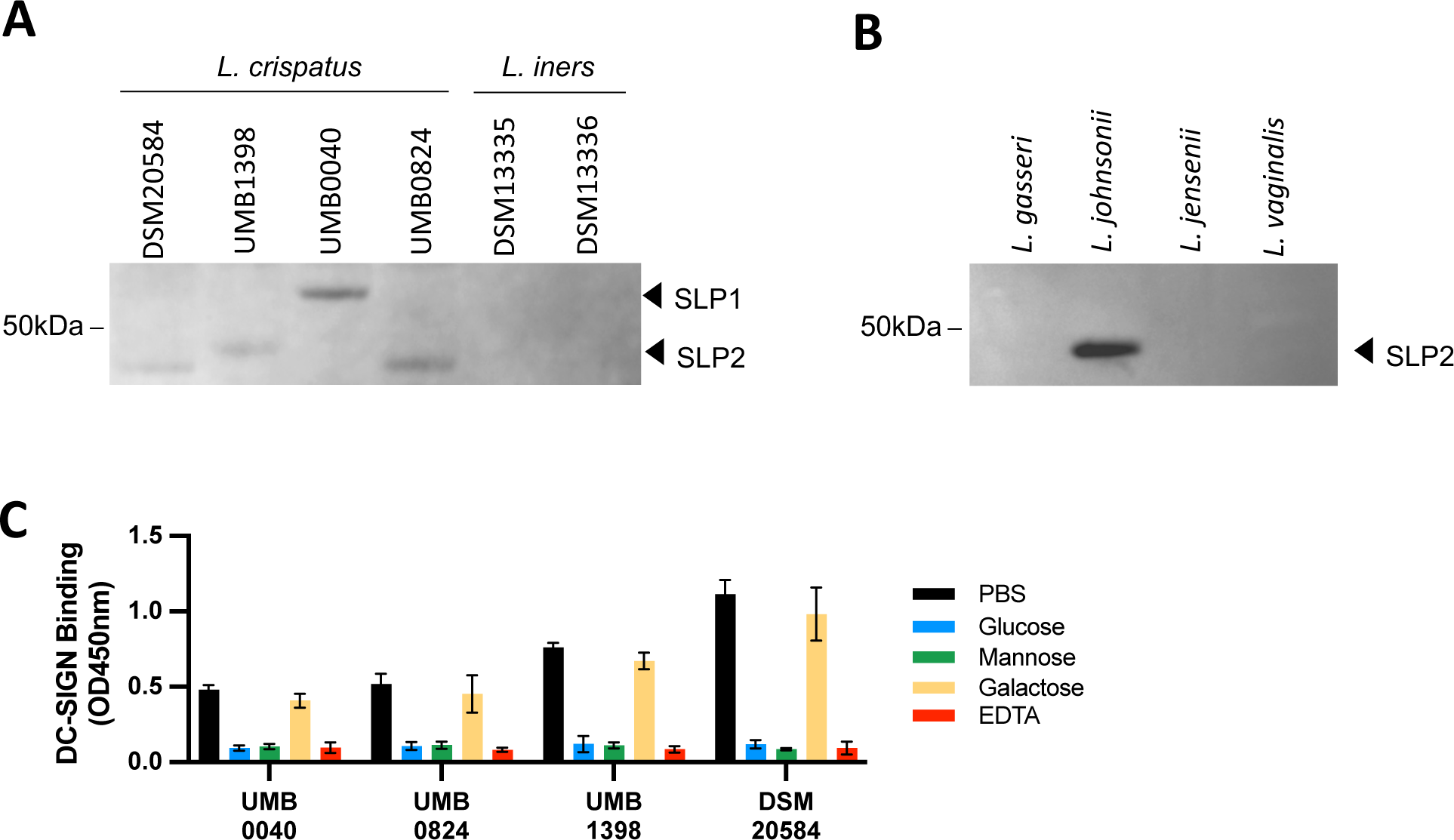
SLPs are ligands of DC-SIGN. (A) Western blotting results for DC-N-Fc binding to LiCl5M extracts from *L. crispatus* and *L. iners* strains (one representative ree replicates), (B) Western blotting results for DC-SIGN-Fc binding to LiCl 5M extracts *L. gasseri* DSM20077, *L. johnsonii* DSM10533, *L. jensenii* DSM20557 and *L. vaginalis* 5487 (one representative of three replicates), (C) Purified SLPs isolated from *L. atus* strains (1ug per well) were coated in 96-well plates and tested for their capacity to DC-SIGN-Fc. DC-SIGN-Fc (1 μg/ml) was pre-incubated or not with 20 mM EDTA or M glucose, mannose and galactose and allowed to react with the SLPs for 2 h at RT. nd proteins were detected using a biotin-conjugated anti-IgG Fc specific antibody and in-HRP, and reading O.D. at 450 nm. Data are mean +/-SD (n=3).

**Extended Data Fig. 5:**
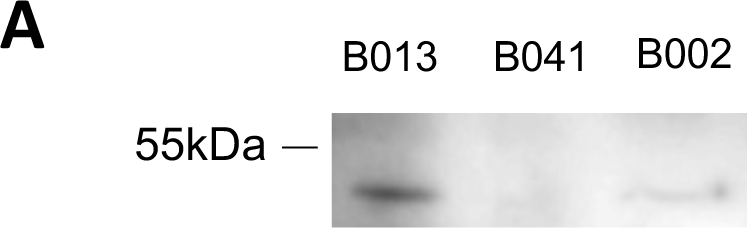
SLPs can be detected in CST-IV dominated CVF samples. Western blotting results for anti-Surface Layer Protein antibody binding to purified CST-IV dominated cervico-vaginal fluid samples (B002 and B013)

## Notes

### Competing Interest Statement

The authors have declared no competing interest.

